# Managed woodlot revealed a trade-off between edible leaves and timber production in *Vitex doniana* Sweet (Lamiaceae)

**DOI:** 10.1101/115154

**Authors:** Sognigbé N’Danikou, Dèdéou A. Tchokponhoué, Aristide C. Houdégbé, Aboègnonhou O.C. Agossou, Enoch G. Achigan-Dako, Françoise Assogba Komlan, Raymond S. Vodouhè, Adam Ahanchédé

## Abstract

*Vitex doniana* Sweet is a major wild-harvested tree resource for food in Benin. However, the species is under threats characterised by increasing human pressure on remnant populations. This study represents the first to explore species’ response to biotic stress. We tested the response of *V. doniana* to coppicing and fertilization. Two stump heights (20 and 40 cm) in combination with three organic manure rates (0.5; 1 and 1.5 kg per seedling), with eight replicates were tested in a randomised complete block design. We used mixed effect models with pseudoreplication, and the maximum likelihood method to compare effects of fixed factors on sprouting vigour, sprout growth and biomass yield in the short (12 months) and medium (5 years) terms. Results indicated that stump height significantly affected sprouting and all growth parameters, in the short and medium terms. However, there seemed a delayed effect of manure. We found initial seedling growth also an important factor. The hidden effect of stump height on biomass yield is discussed. Findings clearly indicate a trade-off between edible leaves and timber production by managed woodlot. Implications of findings for further investigation of above and below ground biomass dynamics and resources allocation in treated trees are discussed.

**Highlights:** A clear trade-off between edible leaves and timber production is observed in managed *Vitex doniana* Sweet woodlot. Coppicing as a biotic stress induced important physiological changes that merit further investigations.

## 1. Introduction

In the optimal foraging theory (OFT) based on the diet breadth model (DBM), domestication represents stress-based prime-mover models in which “*resource depression justifies the engagement of foragers into management as a response to continued pressure for immediate return of high-ranked resources”* (Zeder 2015). Domestication is suggested when there is increased demand of natural products that wild-harvesting cannot meet. Thus, economic demand is thought of to justify the need for domestication (Kupzow 1980). *Vitex doniana* Sweet (a potential tree crop) is an example of wild-harvested species with a sophisticated market chain, which provides tangible revenues that support livelihoods of many households in West Africa, particularly in Benin and Nigeria. However, natural production can no longer meet the ever-increasing demand. As a consequence, human pressure on wild populations has increased (Oumorou *et al.* 2010, Agossou 2011, N’Danikou *et al.* 2011).

While few studies have investigated regeneration of the species and an important body of knowledge is generated (Mapongmetsem 2006, Ahoton *et al.* 2011, Sanoussi *et al.* 2012, Achigan-Dako *et al.* 2014, N’Danikou *et al.* 2014, N’Danikou *et al.* 2015, Mapongmetsem *et al.* 2016), there is a need to explore its horticultural potentials, in order to develop agronomic packages for future cultivation. In this preliminary investigation, we tested the response of *V. doniana* to cultivation management using stump height, initial stump diameter and fertilization (organic manure), three important factors for biomass production in horticultural plants, especially in tree species which are raised for their leaves. In fact, the success of any biomass plantation highly relies on the sustainability of the coppicing and successive pruning system (Hytönen 1994). In this line, a number of both intrinsic and external factors affect stump regeneration among which are stump height, stump diameter, cutting season, cutting method, site quality, fertilization, spacing, rotation length, and species (Hytönen 1994, Saifuddin *et al.* 2010). Knowledge of these factors affecting sprouting ability and biomass yield is necessary to define cutting and harvesting schedules in *V. doniana.* The following hypotheses formed the basis for our study: H1) stump height, initial stump diameter and organic manure are important factors that control regrowth and biomass yield in *V. doniana,* and H2) the applied cultivation treatments to produce edible leaves also affect wood yield in the species.

## 2. Material and methods

### 2.1. Plant material and experimental design

Data were collected on a *Vitex doniana* woodlot raised for 5 years. The trees were obtained from sexually regenerated seedlings at the Agonkanmey’s Research Station of the National Agricultural Research Institute of Benin (INRAB). We tested species’ response to coppicing at different height and organic manure. Two stump heights (C20 = 20 cm, and C40 = 40 cm above ground) in combination with three goat manure rates (D1 = 0.5 kg seedling^-1^; D2 = 1 kg seedling^-1^ and D3 = 1.5 kg seedling^-1^), with eight replicates were tested in a randomised complete block design. In addition to the absolute control (C00), we defined a relative control (C01) where trees received light thinning by removing all leaves from the main stem and branches, but not damaging the apical buds. The combination of factors resulted in eight treatments (Table 1). Seedlings were transplanted at 2 m × 2 m spacing (2,500 stems ha^-1^), and in staggered rows. There were eight blocks in total and each block of 16 m^2^ comprised eight plants. Seedlings were transplanted after they were raised for five months in nursery. Due to the fact that seed germination in *V. doniana* was stochastic at the time of this experiment, seedlings produced were of different ages and heights. To account for this difference in initial growth of transplanted seedlings, we defined height ranges and seedlings with dominant heights were first randomized in the rows, followed by shorter ones. With the limited number of available seedlings, due to the reason given above, the effect of other factors such as planting density and pruning frequency could not be tested in this study. Soil at the experimental site was homogenous and water supply was assumed to be optimal.

**Table 1.**
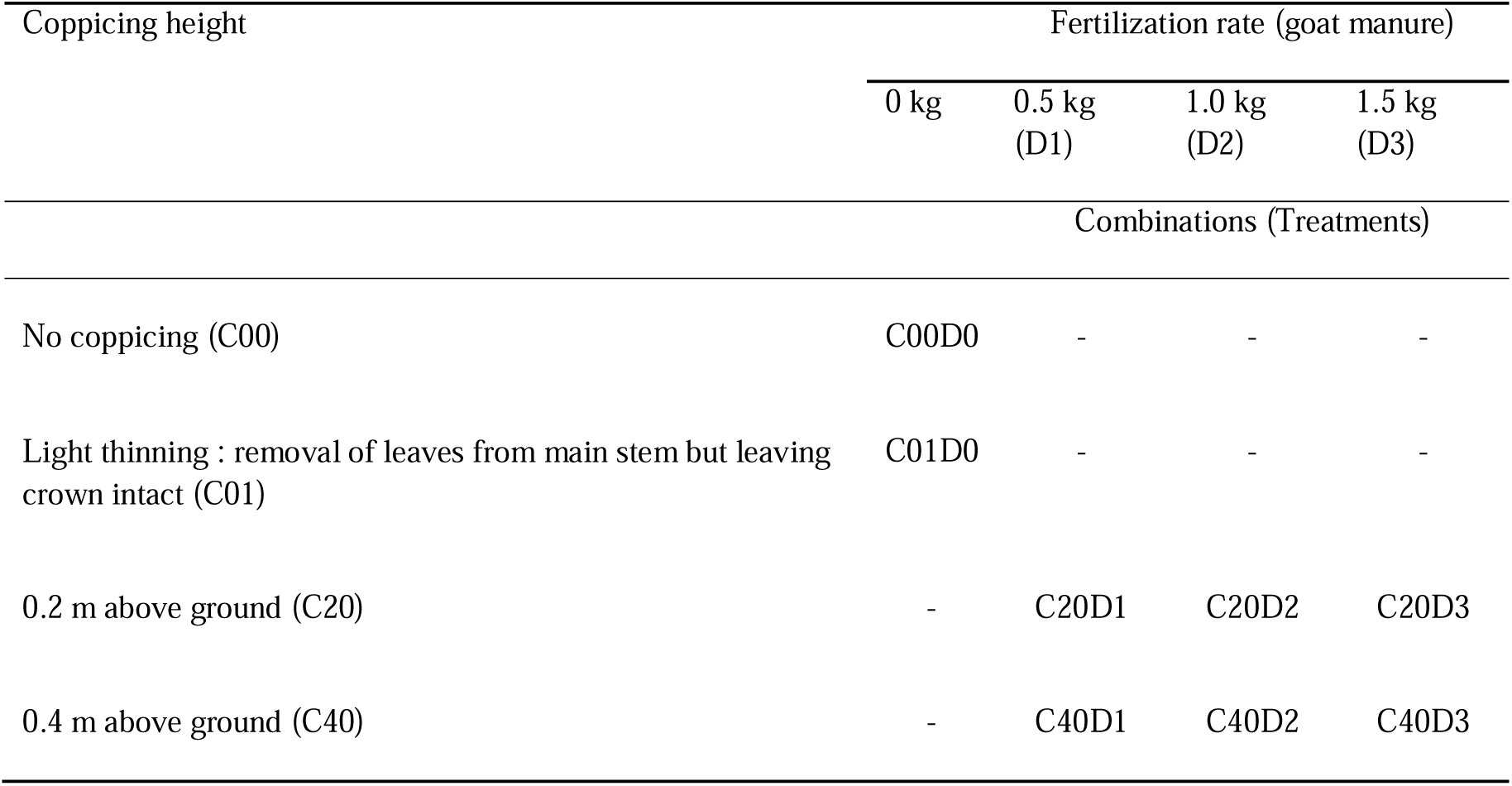
Coppicing and fertilization treatments applied to *Vitex doniana* seedlings

| Coppicing height | Fertilization rate (goat manure) |  |  |  |
| --- | --- | --- | --- | --- |
|  | 0 kg | 0.5 kg<br>(D1) | 1.0 kg<br>(D2) | 1.5 kg<br>(D3) |
| Combinations (Treatments) |  |  |  |  |
| No coppicing (C00) | C00D0 | - | - | - |
| Light thinning : removal of leaves from main stem but leaving crown intact (C01) | C01D0 | - | - | - |
| 0.2 m above ground (C20) | - | C20D1 | C20D2 | C20D3 |
| 0.4 m above ground (C40) | - | C40D1 | C40D2 | C40D3 |

### 2.2. Measurements and data analysis

Height of saplings (from transplanting up to three months), basal stem diameter of trees, number and height of sprouts after each pruning, total wet biomass and dry weight of edible biomass were monitored on temporal basis, over 12 months. Other growth parameters such as basal stem dimeter, diameter at breast height (DBH), number of root suckers, number of branches below 0.2 m, 0.4 m, and below 1.3 m above ground, were recorded when the managed woodlot was three years and five years old.

A total of four prunings were performed on trees that were clear cut at different heights, and six series (every ten days) of measurements of the above parameters were recorded on each tree after each pruning. Pruning was not used as a factor in the current study, but a method to harvest foliage and allow for regeneration. Each series of measurement lasted two months. Thus, we collected 24 series (4 prunings × 6 measurements) of growth data and four biomass yield data over 12 months. DBH was measured only on three years and five years old trees. We used mixed effects models with pseudo-replication to account for temporal autocorrelation across repeated measures on the same trees, and the maximum likelihood method to test the effects of fixed factors (height of pruning and manure application) and random effect (date of measurement, and number of harvest) on the different growth and yield parameters (main stem’s basal diameter and DBH, the number and growth of new shoots, the number of branches below cutting points and below 1.30 m, and the quantity of edible biomass). We also tested effect of the initial basal stem diameter (IBSD-C), measured just before applying the treatments. Selecting the best models that explain the maximum of the variability observed in data, the main effect of IBSD-C was compared to its random effect and this for all response variables. All statistical analyses were performed using R software (The R Core Team 2013).

## 3. Results

### 3.1. Survival of V. doniana to transplantation and to cultivation management

All *V. doniana* seedlings survived after transplantation and also to cultivation management. After three months and before treatments were applied, height of saplings varied between 43 cm to 152 cm (Figure 1). On average, seedlings with bigger initial basal stem diameter at transplanting (IBSD-T) grew better in height than thinner seedlings. Thinner seedlings (IBSD-T ≤ 8 cm) reached 89.00 ± 25.83 cm high, compared with 99.87 ± 24.51 cm for medium size (8 > IBSD-T < 10 cm) and 114.42 ± 17.43 cm for bigger seedlings (IBSD-T ≥ 10 cm). Data also indicated a linear relationship between stem diameter and height growths, in three months old saplings (Figure 2). The variance analysis indicated a very significant effect of initial collar diameter at transplanting on saplings’ growth in height (p < 0.05). Saplings’ height increased with increasing IBSD-T.

**Fig 1.**
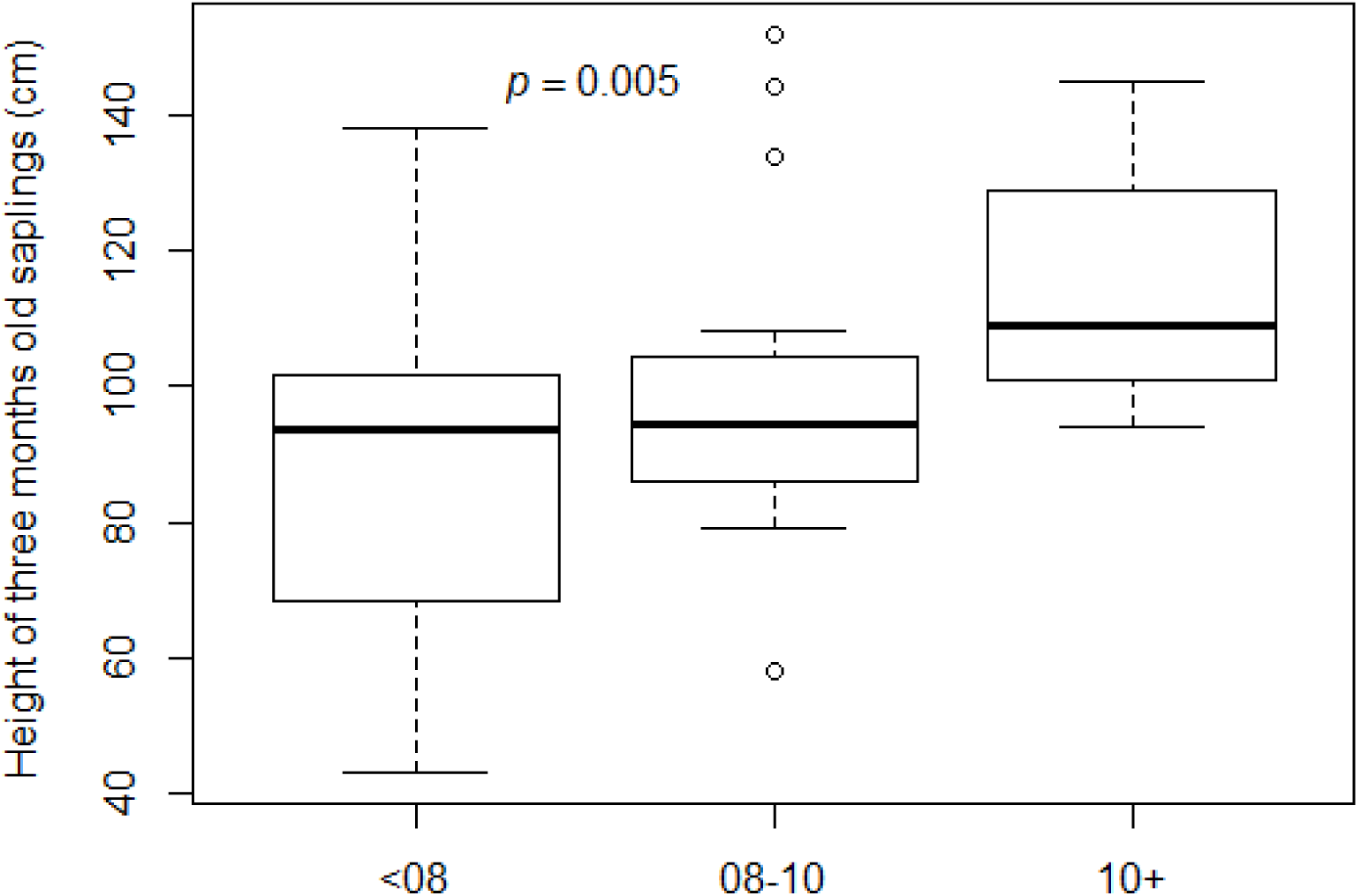
Height of *V. doniana* saplings three months after transplanting and before treatments were applied

**Fig 2.**
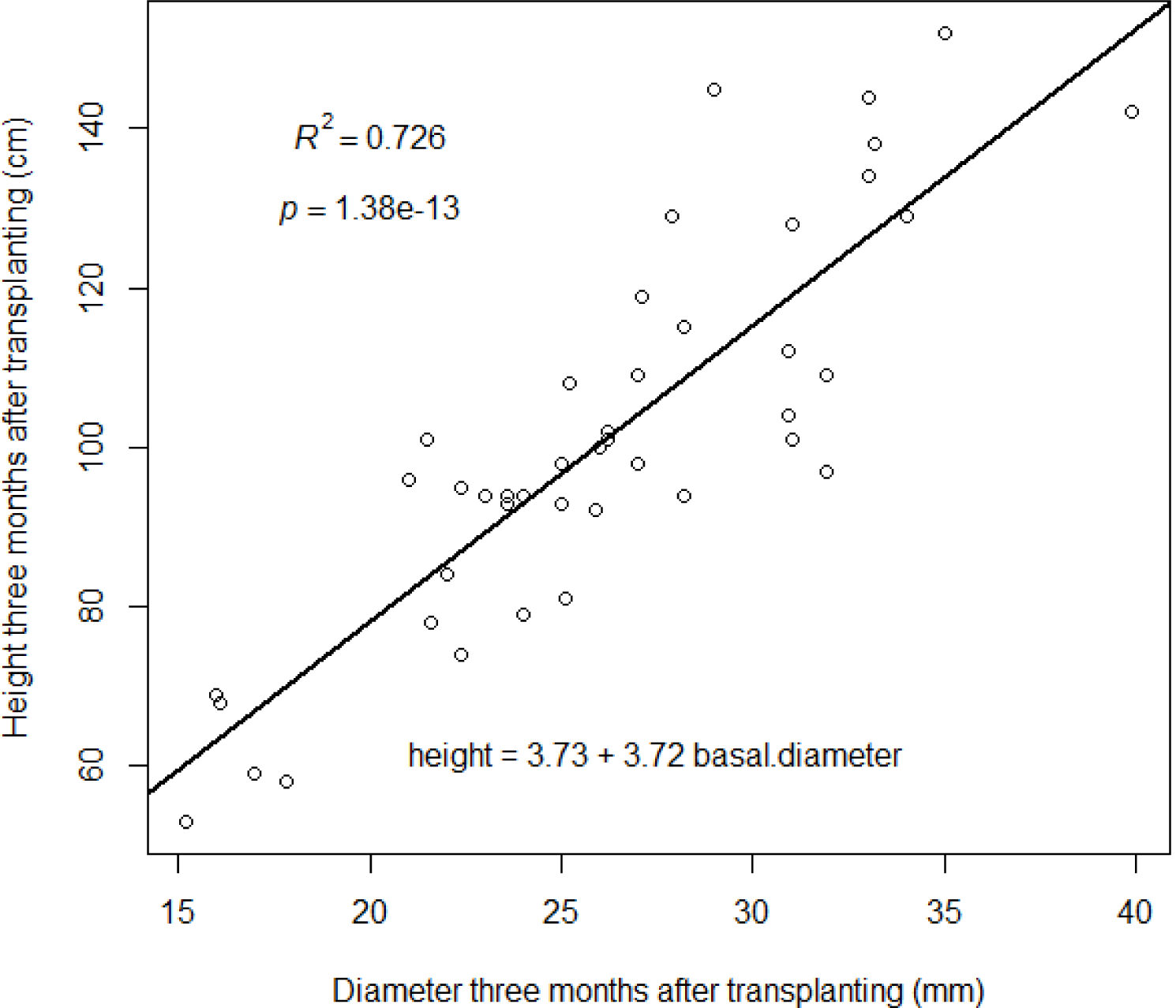
Allometric relationship between stem basal diameter and stem height in young trees

After five years, the survival rate is 100% in both the control trees and in the trees that received cultivation treatments. Data also showed that *V. doniana* is also a fast growing species with the diameter at breast height (DBH) of untreated trees reaching 11.67 cm (6.24 ± 2.60 cm in average) in three years, and 14.65 cm DBH (9.10 ± 3.01 cm in average) in five years old trees. The prediction based on the allometric relationship between height and diameter indicated that under plantation conditions (regular weeding and need-based irrigation) non-coppiced *V. doniana* trees reached confidence interval CI: 2.18 m – 5.89 m (4.14 m in average), and CI: 2.72 m – 7.73 m (5.36 m in average) in three and five years, respectively.

### 3.2. Effects of clear-cutting, manure application and initial stem growth on trees ’ sprouting capacity and biomass production in the short term

#### 3.2.1. Sprouts growth in height

Overall, height of three months old sprouts varied between 20.0 cm and 172.0 cm. The lowest height was obtained in sprouts growing on short stumps while the highest values were recorded on high stumps, all manuring rates considered. On average sprouts from high stumps had higher height compared to those that grew on short stumps, while no linear effect of manure is perceptible (Figure 3). Within each treatment sprouts seemed to show relatively similar height growth after each pruning. The deviance analysis showed that stump height had significant effect on sprouts’ growth in height (p < 0.05), while the effects of manure and IBSD-C were not significant (p > 0.05). Thus, we inferred that after each pruning sprouts height increased with increased stump height.

**Fig 3.**
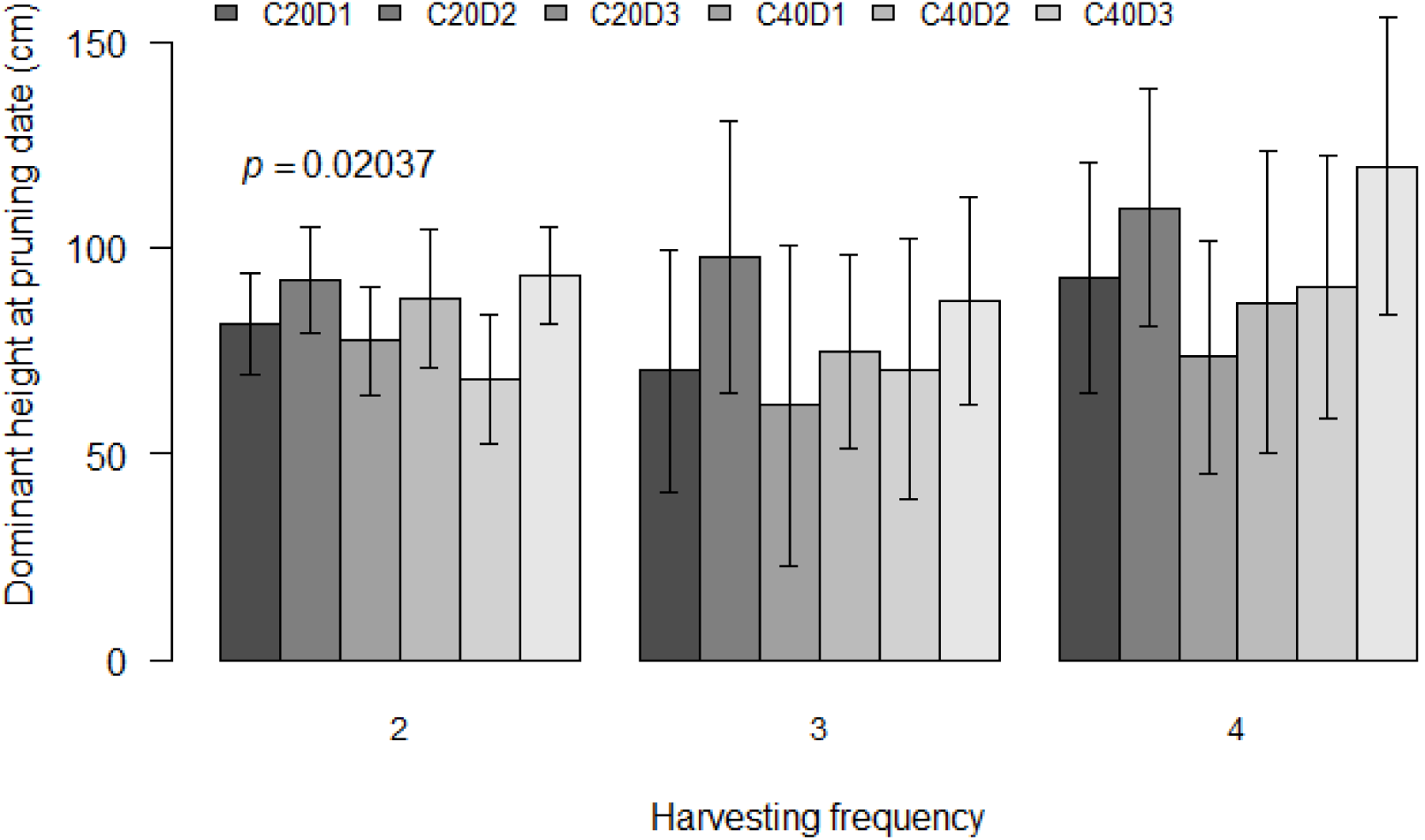
Sprouts growth in height two months after each pruning

#### 3.2.2. Basal stem diameter

Data indicated that irrespective of manure rate, basal stem diameter of mother stand was higher in high stumps (Figure 4). In average, basal stem diameter growth increment ranged from 1.09 mm to 1.14 mm in 20 cm stump height, while it ranged from 1.26 mm to 1.49 mm in trees with 40 cm stump height. It appeared that higher stumps (40 cm above ground) grew better in basal diameter compared to shorter stumps (20 cm above ground), irrespective of manure application. It is also noticed that in average the basal stem diameter growth speed was higher in the first 15 days after pruning (DAP), and then gradually decreased for all treatments, till 45 DAP. From 60 DAP we recorded a new increase in the diameter growth rate. The deviance analysis indicated that stem basal diameter significantly increased with increased pruning height (p < 0.05), while manure application did not show any significant effect (p > 0.05), as shown in Figure 2. There was also no significant interaction effect of stump height and manure application rate on basal stem diameter (p ? 0.05).

**Fig 4.**
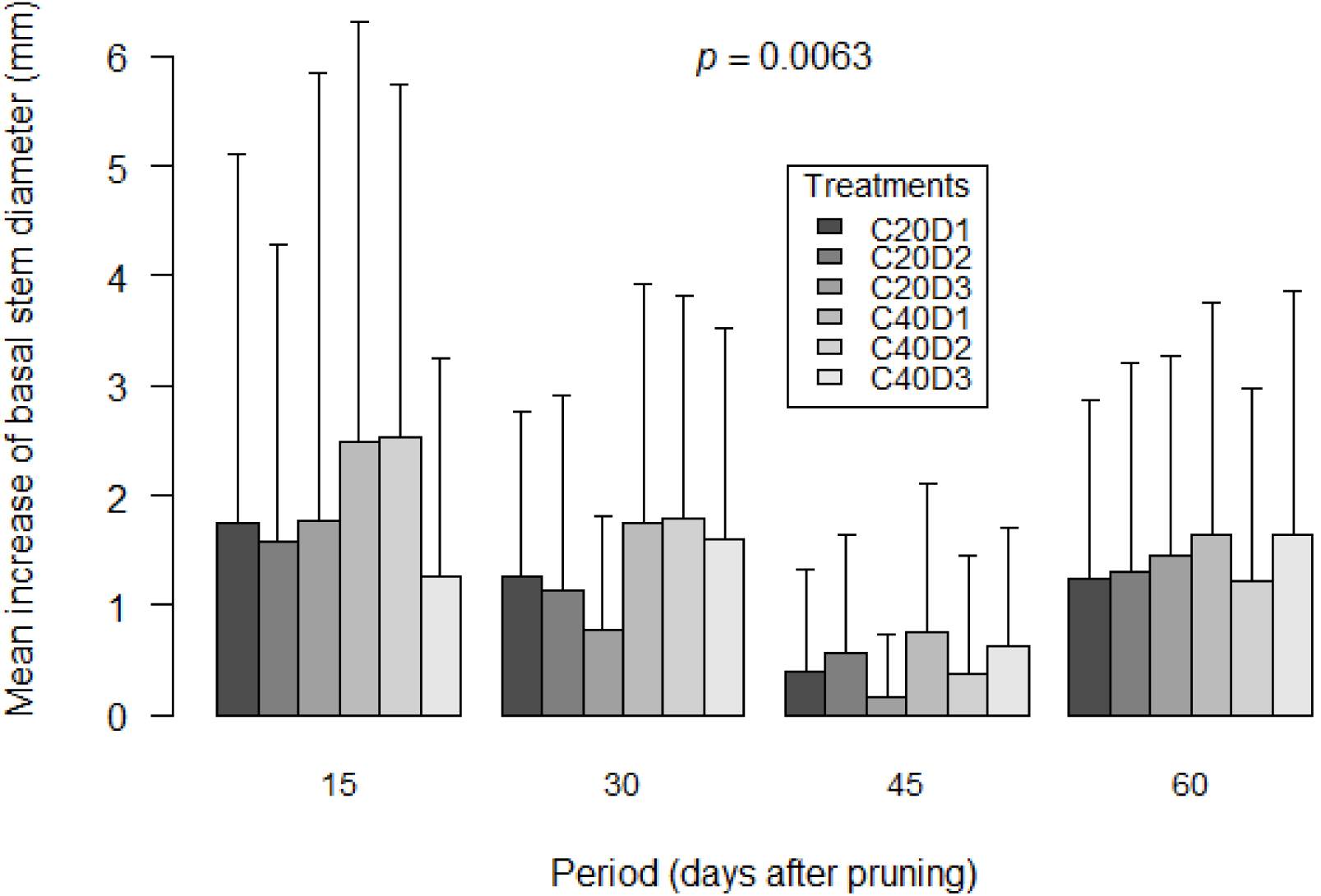
Mean growth rate of basal stem in diameter after clear-cutting and subsequent pruning. Refer to Table 1 for the definition of treatments

#### 3.2.3. Production and growth of sprouts

Results indicated that between two pruning periods, the number of sprouts produced increased over time (Figure 5). The number of sprouts produced after each pruning varied between 0 and 12 (4 sprouts in average). It was observed that while short stumps continued to produce new sprouts for harvest till the 60^th^ DAP (perceptible with C20D1), higher stumps hardly produced new shoots after 30 DAP. The number of shoots produced also seemed to vary with manure rate, with the highest shoots obtained in trees that received 0.50 kg of manure. However, from the graphic there seemed to be no linear effect of manure rate on the number of shoots that were produced, as trees that received higher manure rates (D2 and D3) produced less shoots compared to those which received low manure dosage.

**Fig 5.**
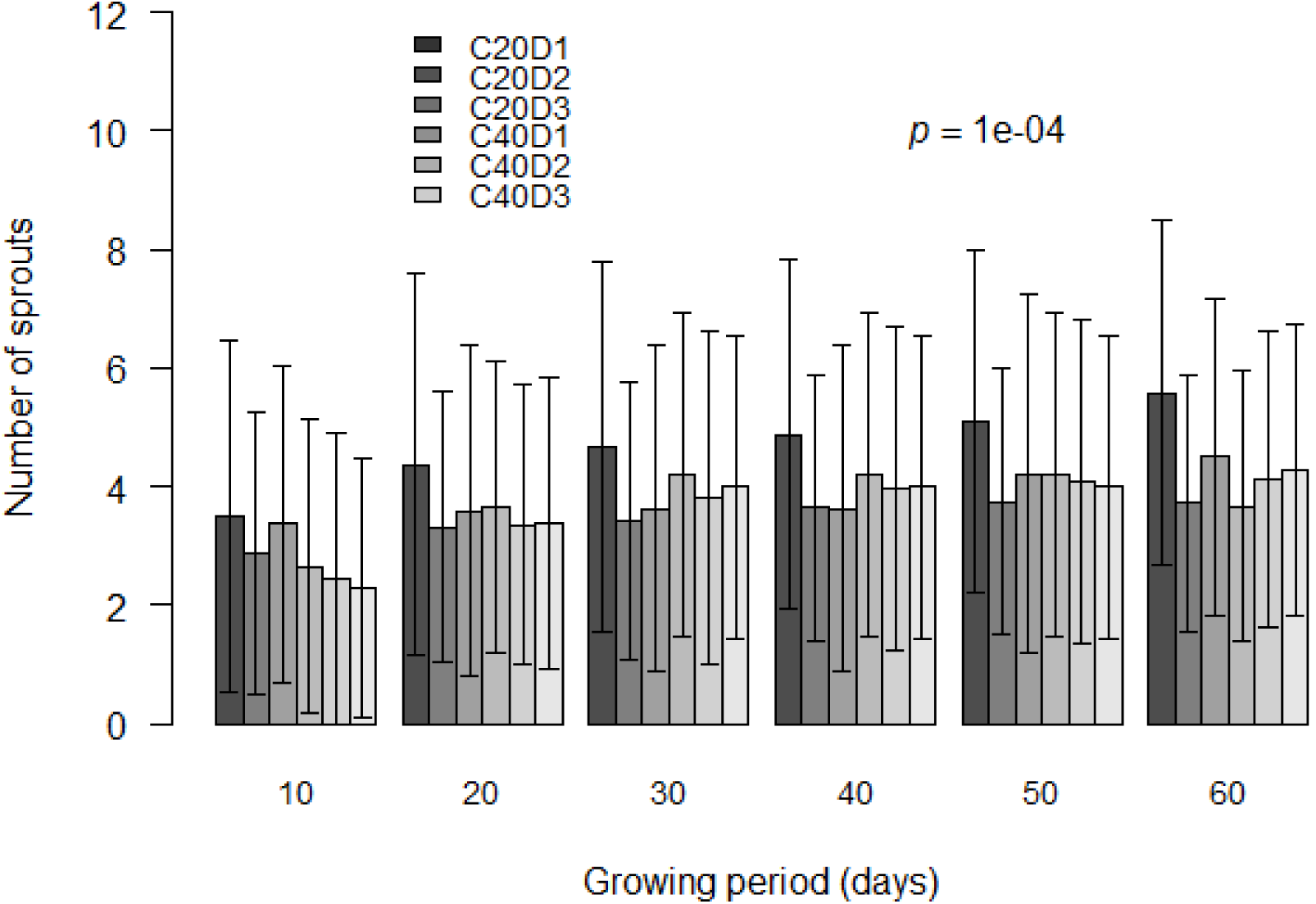
Number of stump sprouts after coppicing, measured periodically.

The deviance analysis indicated significant effects of coppicing (p < 0.05) and IBSD-C (p < 0.001) on the number of sprouts, while the effect of manure application was not significant (p > 0.05). There was a significant two-way interaction effects between stump height and IBSD-C (p < 0.001), and between ISBD and manure (p < 0.001) on the number of sprouts, while the interaction between stump height and manure was not significant (p > 0.05). The three-way interaction effect between factors was also significant (p < 0.001). It is therefore concluded that the number of sprouts increased with the increase of initial basal stem diameter before clear-cutting (Table 2), while it decreased with stump height. Short stumps with IBSD-C ≥ 3 cm produced higher number of sprouts, even with low manure rate.

**Table 2.** Mean number of sprouts per initial basal stem diameter class before clear-cutting (IBSD-C)

| IBSD-C classes<br>(mm) | Days after pruning |  |  |  |  |  |
| --- | --- | --- | --- | --- | --- | --- |
|  | 10 | 20 | 30 | 40 | 50 | 60 |
| 10-25 | $2.47 \pm 2.39$ | $3.26 \pm 2.26$ | $3.45 \pm 2.4$ | $3.64 \pm 2.32$ | $3.71 \pm 2.3$ | $3.76 \pm 2.13$ |
| 25-30 | $3.00 \pm 2.31$ | $3.69 \pm 2.71$ | $4.20 \pm 2.97$ | $4.23 \pm 2.97$ | $4.44 \pm 2.97$ | $4.42 \pm 2.77$ |
| $\geq 30$ | $3.23 \pm 2.97$ | $4.02 \pm 2.90$ | $4.38 \pm 2.79$ | $4.44 \pm 2.72$ | $4.73 \pm 2.83$ | $5.00 \pm 2.71$ |

Within 60 days twigs length varied between 0.40 cm and 104.80 cm, when considering all treatments (Figure 6). Overall, the mean length was higher in all twigs that sprouted on high stumps, irrespective of the manure rate. The deviance analysis indicated that among the tested factors, only stump height had significant effect on twigs growth (p < 0.05), while no significant effects of goat manure and IBSD-C were observed (p > 0.05). There were significant 2-way interaction effects between stump height and manure, stump height and IBSD, and between manure and IBSD-C (p < 0.05, in all cases). We concluded that twigs growth increased with increased stump height. On short stumps, sprouts grew faster when IBSD-C is small (<25 cm), while on higher stumps, sprouts grew faster when IBSD-C is big (≥ 30 cm).

**Fig 6.**
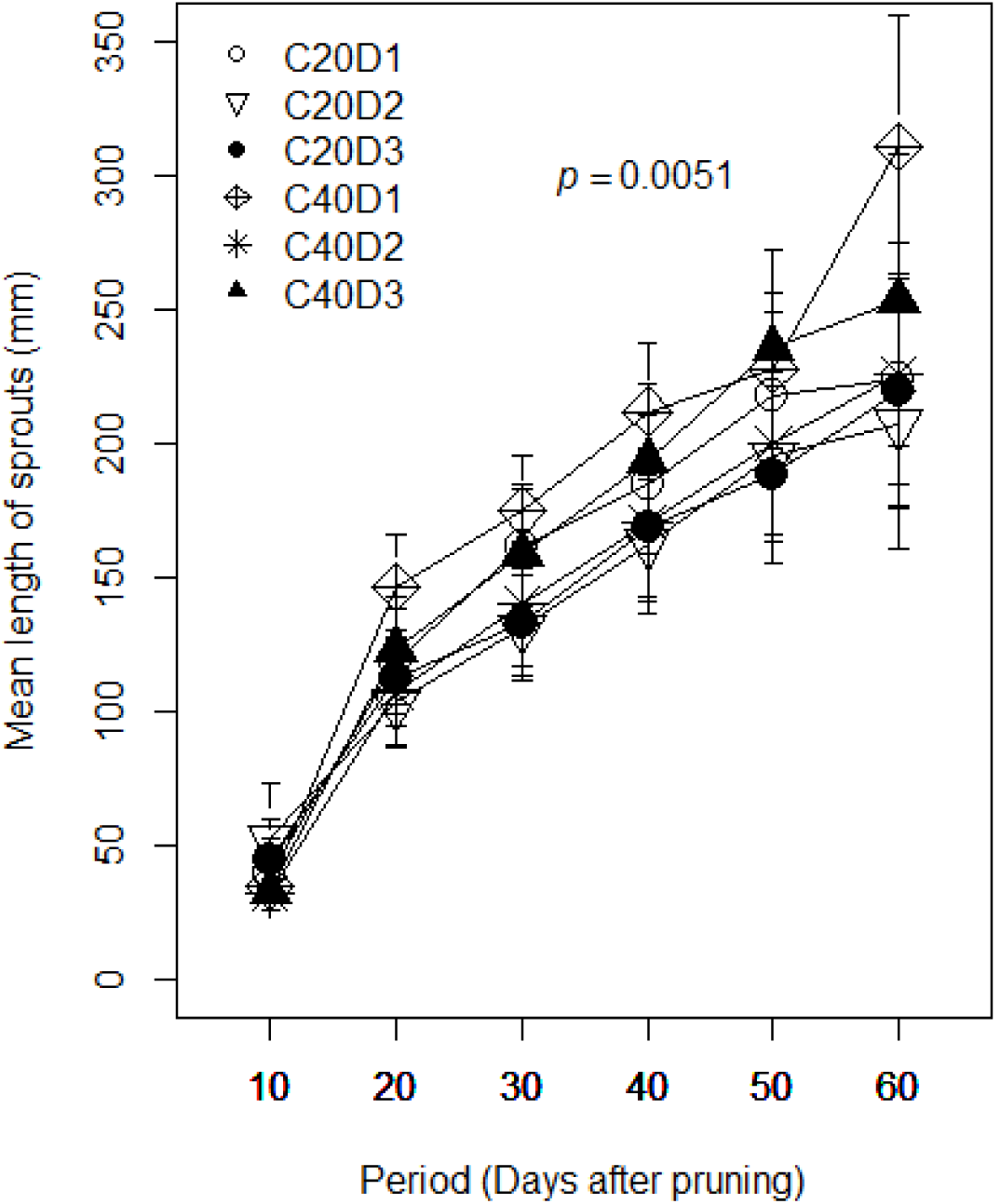
Growth in twigs length per stump height and manure rate, over 60 days after pruning. Refer to Table 1 for the definition of the treatments.

#### 3.2.4. Edible biomass

Overall, dry matter varied from 0.98 g to 12.98 g per plant, equivalent to 2.45 kg to 32.45 kg per hectare per harvest. The highest yield was obtained in the first harvest. *Vitex doniana* leaves contained in average 32% dry matter per 100 g of edible portion. The quantity of edible dry matter also decreased over time, to become very low after three harvests (Figure 7). In the first harvest after clear-cutting, biomass produced by short stumps, regardless the amount of manure applied, was slightly higher than that by high stumps. However, the reverse was observed after the second and third harvests where biomass produced by shorter stumps was lower compared with that measured in high stumps, this irrespective of the manure dosage. Biomass yield in short stumps was 8.8% less in the second and 127% less in the third harvest, respectively. It’s worth indicating that the last harvest coincided with the period of lowest rainfall in the year (January). Nonetheless, deviance analysis indicated there is no significant effect of coppicing and manure application on the dry matter produced (p > 0.05). In addition, a very significant difference was noted in the amount of dry matter over a period of three harvests (p < 0.001). Biomass decreased with increased harvest. It was also found a marginal interaction effect between stump height and number of harvests on biomass yield (p = 0.07).

**Fig 7.**
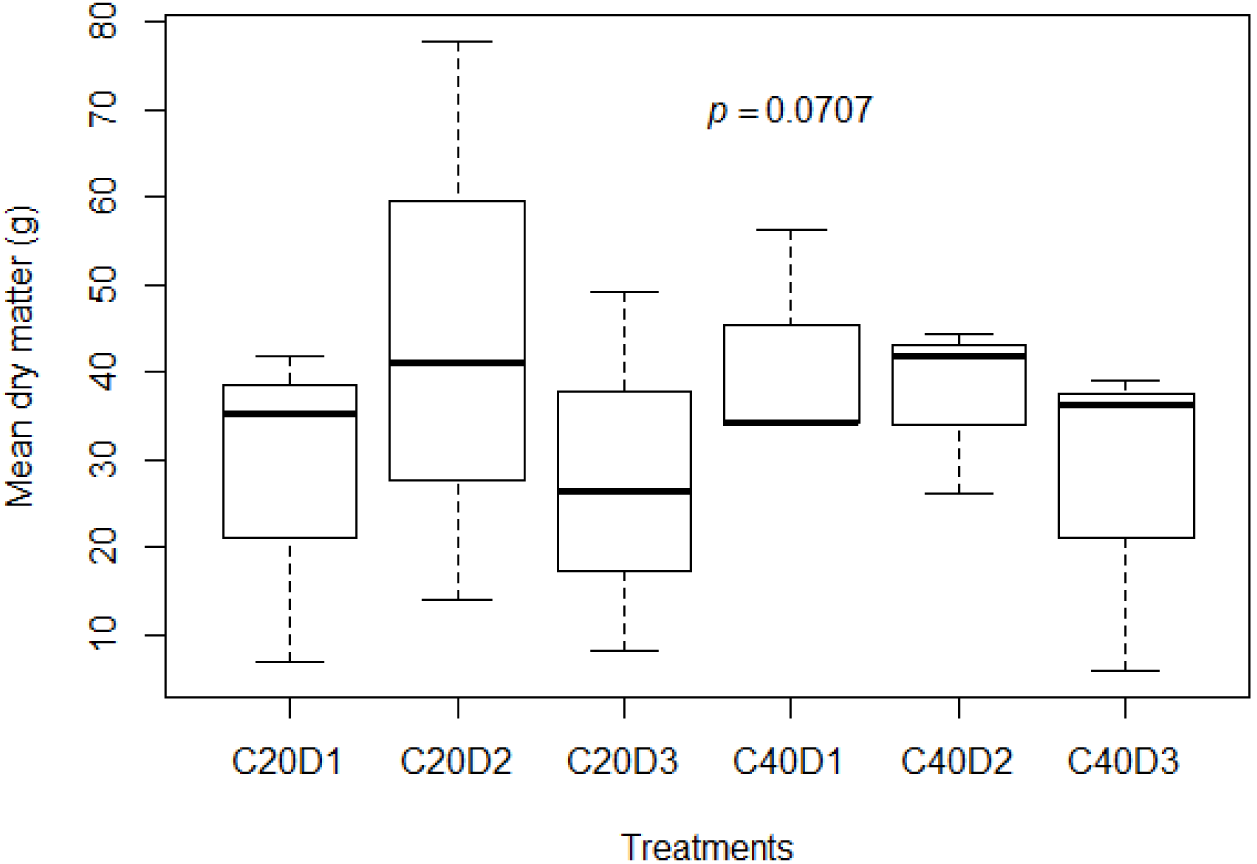
Mean dry matter of edible biomass per treatment over the whole experiment

### 3.3. Effects of cultivation management in the medium term

#### 3.3.1. Diameter at breast height

Overall, the DBH ranged from 0.66 cm to 11.67 cm in three years old trees, and 0.95 cm to 14.65 cm in five years old trees. In average, the DBH was higher in the control (6.24 ± 2.60 cm and 9.10 ± 3.01 cm in three and five years old trees, respectively) (Figure 8). Smaller DBH were recorded in sprouts that grew from short sumps (3.00 ± 1.91 cm and 4.68 ± 2.45 cm in three years and five years old trees, respectively). The deviance analysis showed that only stump height significantly affected DBH (p < 0.001), while the main effects of manure (p > 0.05), and initial basal diameter before clear-cutting (p = 0.08) on DBH were not significant. There were significant two-way interaction effects between stump height and manure, and between manure and IBSD-C (p < 0.05, in all cases). However, the interaction between stump height and IBSD-C was not significant (p = 0.07). We concluded that DBH increased with increased stump height and with decreased manure rate, and also with increased IBSD. It was also found that DBH of trees that received light thinning (C01D0) was not significantly different from the absolute control (p > 0.05).

**Fig 8.**
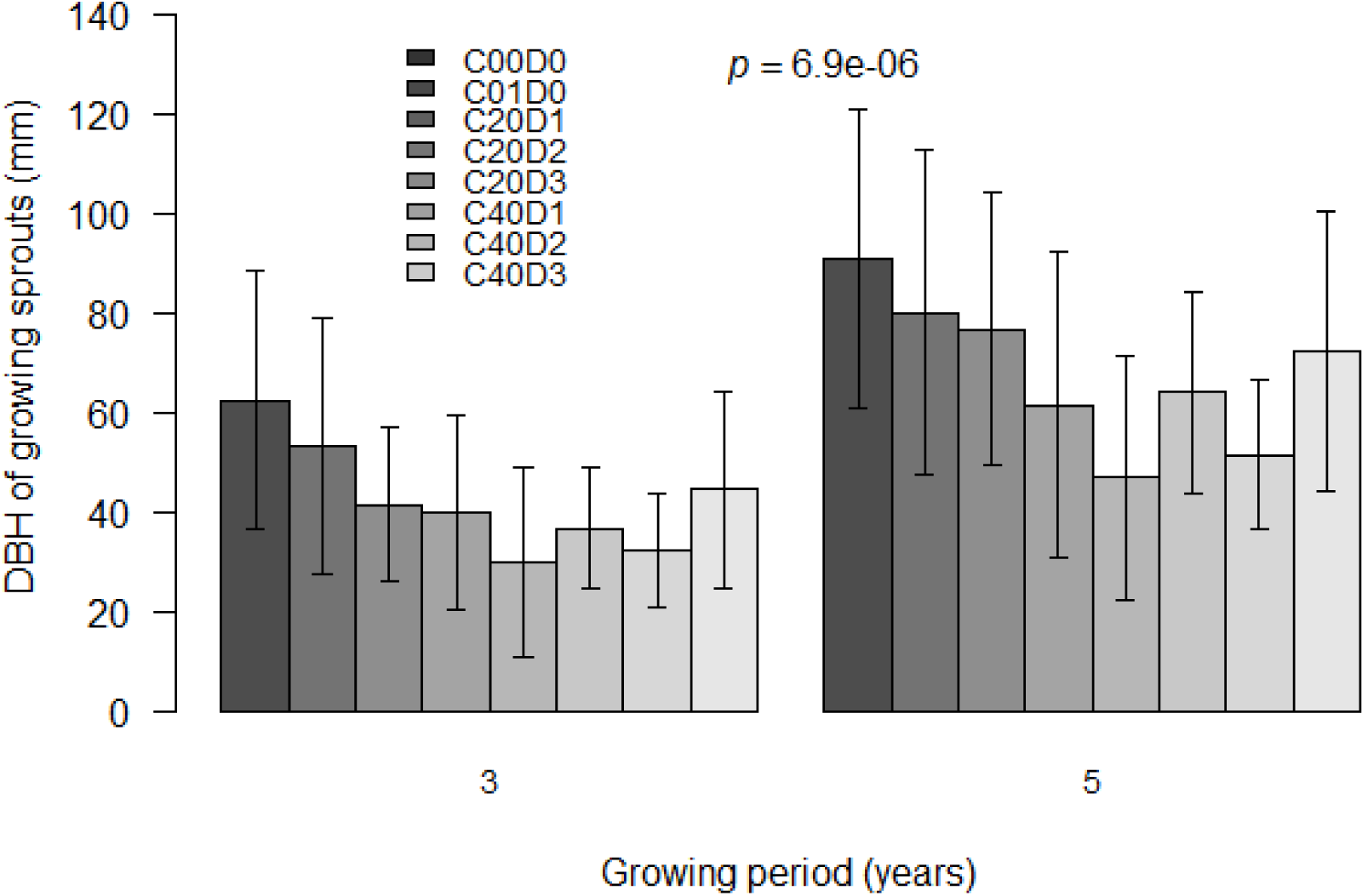
Diameter at breast height (DBH) of 3 and 5 years old *V. doniana* trees

#### 3.3.2. Basal stem diameter

Stem basal diameter of the absolute control trees (D0C00) was in average 11.05 ± 2.36 cm and 14.33 ± 3.18 cm, for three years old and five years old trees, respectively. Trees that received light thinning (C01) showed a slightly lower stem growth; 10.21 ± 2.17 cm and 13.47 ± 4.19 cm in three years and five years old trees, respectively. Overall, slower diameter growth was observed in stumped trees which showed lower basal stem diameter (Figure 9). Deviance analysis indicated a very significant effect of stump height and IBSD-C on basal stem diameter after five years (p < 0.01, in both cases), while the effect of manure was not significant (p > 0.05). There were also significant two-way interaction effects between stump height and manure, stump height and IBSD-C, and between manure and IBSD-C (p < 0.05, in all cases), while no significant three-way interaction effect was observed. We inferred that basal stem diameter increased with increased stump height and with decreased manure rate. Overall, stumps with big IBSD-C relatively grew faster.

**Fig 9.**
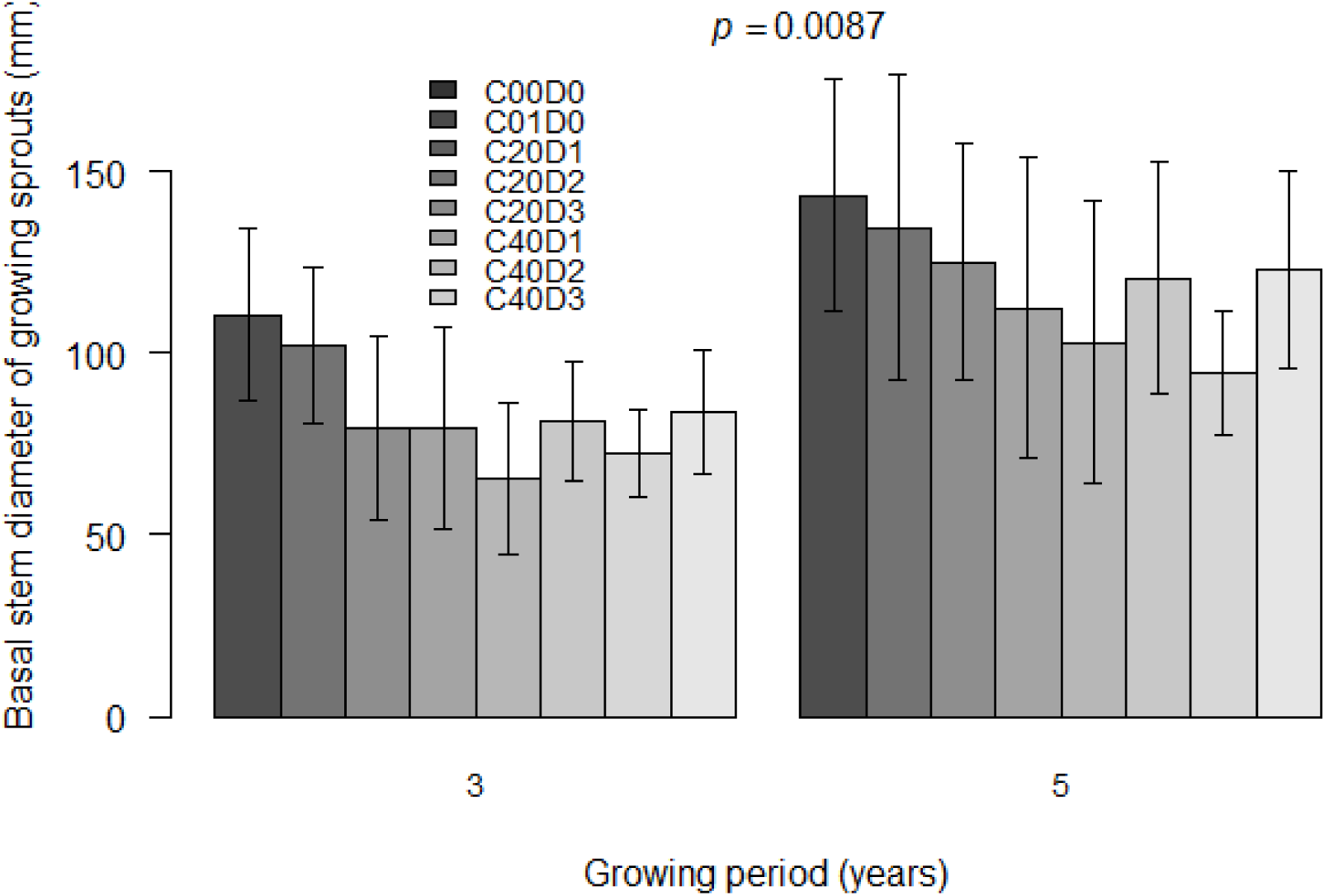
Average basal stem diameter in three and five years old trees, per treatment

#### 3.3.3. Production of branches

Overall the number of branches on *V. doniana* trees increased from the ground to the top of tree (Figure 7). The trend indicated that the trees which received cultivation management treatments continued to produce relatively higher number of branches below cutting points (Figure 10 A and B) and below 1.30 m (Figure 10 C), irrespective of their initial basal stem diameter before clear-cutting. Overall, the number of branches in total increased from three years old to five years old. However, there were more branches below 1.30 m and below cutting points in three years old sprouts compared to in five years old ones. The deviance analysis indicated that three years after treatments were applied there is still a significant effect of stump height on the number of branches below the cutting points of 0.20 m and 0.40 m, and on the total number of branches (p < 0.05, in all cases). The effect of IBSD-C was significant only on the number of branches below 0.20 m (p < 0.05). The main effect of manure application was not statistically significant in any case. Nonetheless, there were significant two-way interaction effects between stump height and manure, stump height and IBSD-C, and between manure and IBSD-C, though only on branching below 0.20 m (p < 0.05, in all cases). Results therefore indicated that the number of branches produced by three years old trees increased with decreased stump height and with increased manure rate. The number of branches also increased with increased IBSD-C.

**Fig 10.**
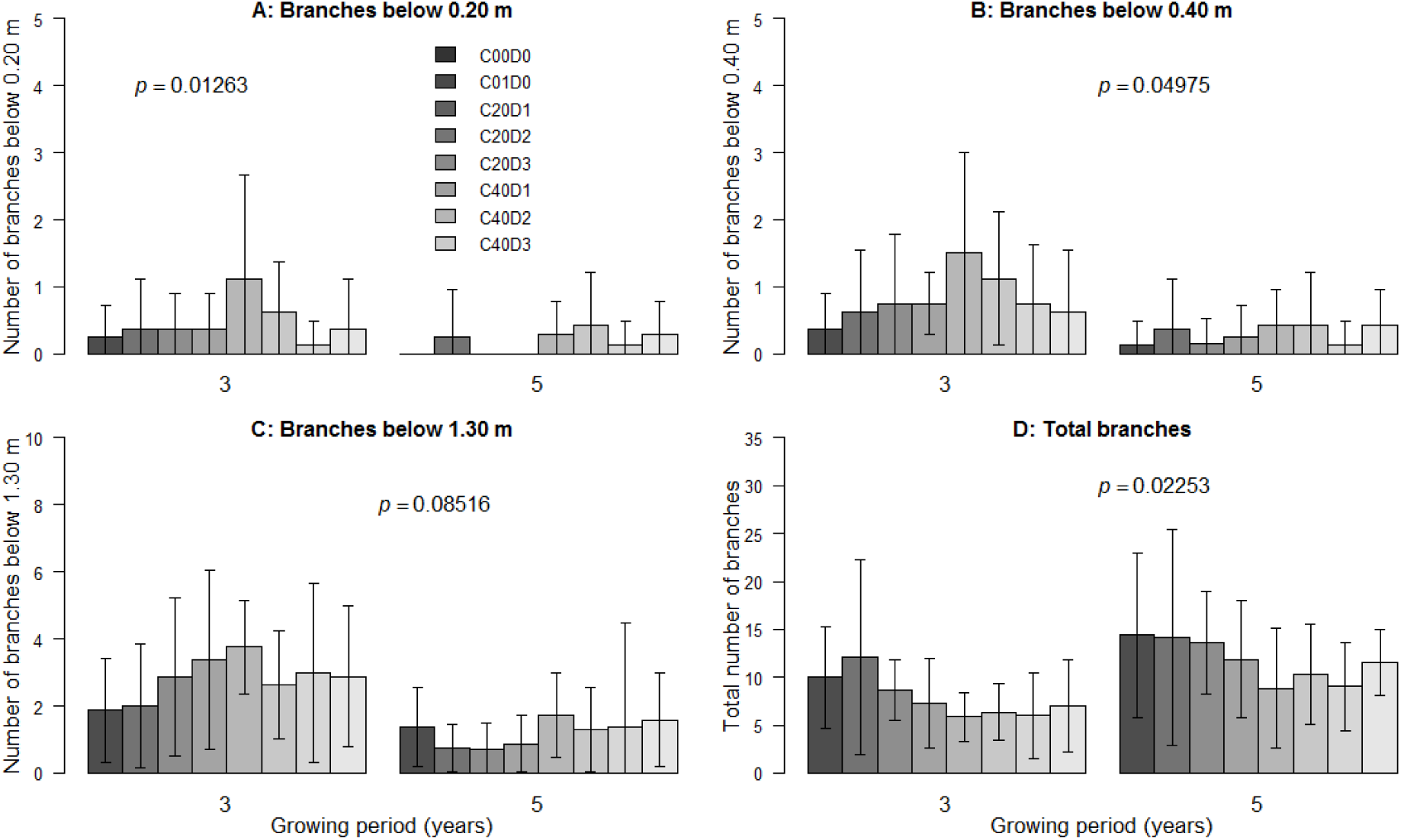
Mean number of branches per tree over three and five years of woodlot management

#### 3.3.4. Production of root suckers

The average number of root suckers produced by three years old trees was higher in control treatments compared with coppiced and fertilized trees (Figure 11). It is also observed that the average number of root suckers decreased in five years old trees. High stumps with big IBSD-C irrespective of manure rate produced relatively higher number of root suckers. The deviance analysis indicated a significant effect of stump height on the number of root suckers produced by three years old trees (p < 0.05), while the effects of manure and IBSD-C were not significant (p > 0.05). The interaction effects were not significant. We therefore concluded that the number of root suckers increased with increased stump height.

**Fig 11.**
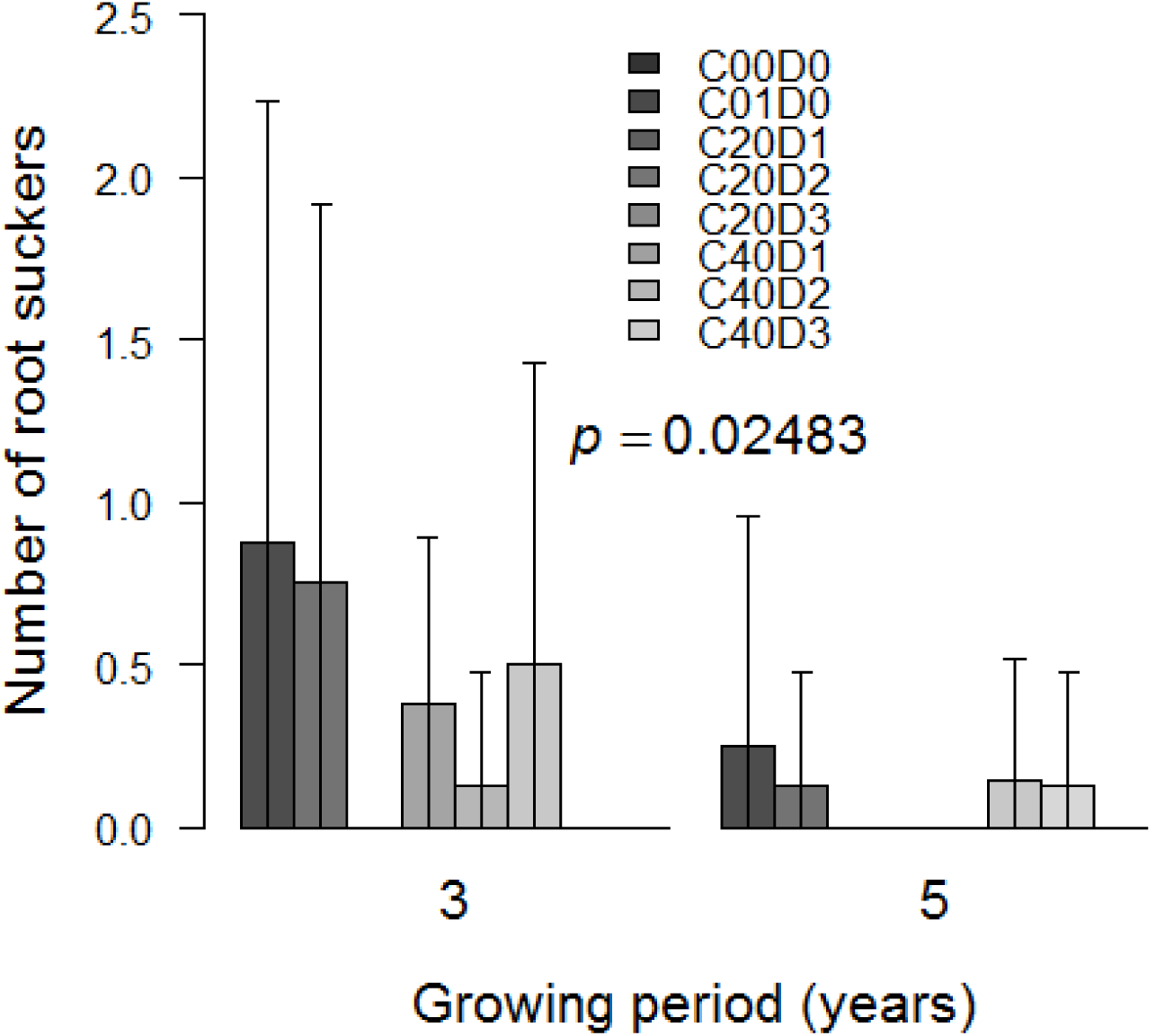
Average number of root suckers produced per tree per treatment, by three years and five years old trees

## 4. Discussion

### 4.1 Growth and development

Our study represents the first to investigate *V. doniana* growth and development in plantation. The findings indicated that it is a fast growing species, with height of non-coppiced trees ranging between CI: 2.17-5.89 m (4.14 m in average) three years after transplantation. Five years plantation trees reached between CI: 2.72-7.73 m high (5.36 in average). These values are by far superior to the 1.70 m reported by Ky (2008), though this latter value likely referred to wild populations. DBH of untreated trees ranged between CI: 2.60-11.7 cm (average 6.24 cm) and CI: 4.46-14.65 cm (average 9.10 cm) in three and five years, respectively. These values of DBH are comparable to that of *Tectona grandis* Lf, another commercial timber species of the Lamiaceae family, where mean quadratic circumference of young plantation trees (below 5 years) was 23,13 ± 0,62 cm (ca. 5.43 cm DBH), in Southern Benin (Atindogbe *et al.* 2012). Most interesting, non-coppiced five years *V. doniana* trees showed higher DBH compared with the 29,67 ± 1,21 cm circumference (ca. 6.15 cm DBH) of high teak plantation forest of more than five years old (Atindogbe *et al.* 2012). Taking into account that *V. doniana* timber has physical properties closely comparable to that of *T. grandis,* (Makonda *et al.* 2016) the former represents a potential commercial multi-purpose tree crop.

### 4.2 Coppicing to promote sprouting, sprout growth and biomass production

The main product collected on *V. doniana* is its young leaves used for vegetable. Therefore, the effects of agronomic techniques on leaves production in horticultural systems are important to investigate. Our results indicated that stump height is crucial for sprouting intensity, with short stumps producing more sprouts than higher ones. This response was reported in some species, and was partly explained by the fact that short stumps are more likely to be connected with the roots which increases the availability of water and metabolites that are needed in sprouting-buds (Hytönen 1994, Wilson 1994, Thomas and Schiefelbein 2004). It was also reported that removal of leaf area significantly affects the ability of sprouting, sprouts growth and root formation (Wilson 1988, Wilson 1994, Paukkonen and Kauppi 1998, Thomas and Schiefelbein 2004, Dharmakeerthi *et al.* 2008, Saifuddin *et al.* 2010). Partial or total removal of the above-ground biomass of trees can result in interruption of root growth or partial death in the root system (Berninger and Salas 2003, Dharmakeerthi *et al.* 2008). Leaves removal is assumed to be detrimental to root formation because they supply root promoters such as indole 3-acetic acid (IAA), auxin and other rooting cofactors, vitamins, carbohydrates, organic nitrogen (Wilson 1988, Hartmann *et al.* 2011). Thus, greater the leaf area, the better the root and shoot growth. This explains that leafy stumps and cuttings are likely to promote more roots and shoot growth than leafless stumps. In addition, high (leafy) stumps had larger number of buds and thus stronger sink activity which would therefore increase the metabolic activity and health of leaves. Leaves will then continue to supply photosynthates and rooting cofactors (Thomas and Schiefelbein 2004). Our findings are consistent with this observation where (leafless) 20 cm stumps, although producing higher number of sprouts, did not result in better growth compared with higher (more leafy) stumps. In our experiment coppicing should have induced a slow growth in root biomass, and with more severe effect on short stumps’ root system development. The sprouts which developed on stumps with more vigorous root system (40 cm stumps) will, over time, likely grow faster compared with sprouts that developed on stumps with poor root growth (20 cm stumps). The same trend was described for walnut tree (Balandier *et al.* 2000) and *Eucalyptus* (Wilson 1988, Wilson 1994), where increased reduction of leaf area reduced rooting and sprouts growth. This is in agreement with theories on balanced growth in plants (Davidson 1969). It means that coppicing or pruning modifies the shoot-root ratio and the response of affected trees is either the quick activation of sprout-producing buds to trigger the photosynthetic activity at the expense of other organs such as roots (case of leafless short stumps) or, when at all possible, to prioritize the supply of rooting cofactors and carbohydrates to sustain roots’ function (case of leafy high stumps). The effect of stump height on sprouting is also partially attributed to the location of sprout-producing buds on the main stem (Hytönen 1994). It is observed in *V. doniana* that the number of buds increased with the stem segment considered. The last bottom segment of stem (close to ground) has more sprout-producing buds than upper segments. In line with the above, the low number of root suckers observed in stumped trees can be explained by the reduced growth of their root systems.

In the majority of cases reported, biomass increased with increased stump and pruning height (Yasui and Fujie 1971, Tipu *et al.* 2006, Saifuddin *et al.* 2010). Nonetheless, although rare, the opposite was also reported in some species (Hytönen 1994). Conceptually, biomass yield is known to be affected by the amount of available carbohydrates for the growing trees (Beminger *et al.* 2000,Berninger and Salas 2003). Therefore, the most plausible scenario in our study would be that more leafy stumps (here high stumps) would produce higher biomass compared with leafless stumps (short stumps). However, the effect of stump height on biomass may not be conclusive after only one rotation experiment. Our results indicated that biomass was higher in short stumps at the first harvest after coppicing, and then the trend changed in the subsequent harvests, where high stumps now produced higher biomass throughout the rest of experiment. Also, the data reported in this study represents only one rotation and this could explain our finding that there was no significant effect of stump height on dry biomass for the first rotation. The season of coppicing also affects biomass production by coppiced trees in many species, with the dormant season being more favourable for the harvest of protein-rich leaves (Hytönen 1994). Decline in biomass production within a rotation can also be due to the frequency of harvest. High harvesting frequency also led to rapid decrease in biomass production (Hytönen and Issakainen 2001, Saifuddin *et al.* 2010). In our study, *V. doniana* was harvested every two months and could explain the rapid decline of biomass yield from one harvest to the next. Although seasonality was not assessed in the current trial, the decline in biomass (dry matter) yield could also be due to the season in which trees were harvested. In fact, the last harvest happened in the driest month and reduced water availability could explain the low yield.

### 4.3 Ambiguous effect of organic manure on growth and yield of indigenous trees

Response of indigenous tree species to organic manure varied with species and soil quality. While a beneficial effect of poultry manure on growth and yield parameters was noted in some plantation species such as *Theobroma cacao* L. (Orozco and Thienhaus 1997), sole application of poultry and farmyard manure had no significant effect on Balady mandarin tree *(Citrus reticulata* Blanco) where an increase of manure rates decreased all growth and yield parameters (Gamal and Ragab 2003). Also, no growth and yield difference was obtained with increased fresh and processed poultry manure application on fruit bearing and non-bearing citrus trees (Ferguson 1994). Besides, some indigenous species showed no response to either organic or inorganic manure. This is the case of *Uapaca kirkiana* Müll.Arg. and *Sclerocarya birrea* (A.Rich.) Hochst. where application of compost, chemical fertilizer and dry-season irrigation did not increase early growth and survival of plants (Akinnifesi *et al.* 2008). In our study, an increase of goat manure rate did not improve sprouting nor growth and yield parameters in *V. doniana,* in the short term. Within the first 12 months we observed that stumps that received 0.5 kg of manure produced relatively higher number of sprouts compared with those that received 1 kg and 1.5 kg of manure, though the difference was not statistically significant. This negative effect of organic manure on growth parameters was reported on *T. cacao* where even in combination with inorganic fertilizer, a ratio above 50% organic manure decreased growth and yield (Gamal and Ragab 2003). Best survival and growth was also observed in *S. birrea* trees that did not received any management treatments (Akinnifesi *et al.* 2008). This corroborates our finding where trees of the control treatments showed better stem growth (basal stem diameter and DBH). Nonetheless, the fact that in three and five years old trees, though not statistically significant, the number of branches relatively increased with increased manure rate could indicate a delayed uptake of organic manure by the species. Overall, our hypothesis 1 is partially rejected and we conclude that organic manure may not be a factor that control regrowth of coppiced trees.

### 4.4 Trade-off between leaves and timber production by managed V. doniana woodlot

Findings indicated that coppicing and subsequent pruning allowed for more sprouts development which can be harvested for consumption, in the short but also in the medium term. This confirms partially our hypothesis 1 that stump height is an important factor for growth and biomass yield in *V. doniana.* In fact, coppiced trees continued to produce more branches at lower height (below 0.40 m) compared with control treatments. This impact of coppicing on plant’s architecture represents an advantage for horticulture prospects, because *V. doniana* is a big tree when fully developed. In the meantime, it was also observed that coppiced trees to produce edible leaves had the lowest DBH, compared with control treatments. This finding confirms our hypothesis 2 that the applied cultivation treatments to produce edible leaves affect wood yield in the species. The drastic reduction of stem growth in those trees intensively harvested for their leaves clearly indicates a trade-off that should be taken into account in production objectives. Simulation studies using growth models have showed that frequent pruning lead to depletion of reserve carbohydrates in the stems (Berninger *et al.* 2000). Therefore, leaves will be produced at the expense of stem growth. This result implies that natural stands that are intensively harvested in the nature are subject to decline, due to reduced growth which will affect their reproductive capacity. However, though coppicing affected timber stock, the coppiced and frequently pruned trees can yield a substantial quantity of firewood, which was not measured in the current study.

### 4.5 Implications for research

Our study showed contrasting effects of stump height on sprouting vigour and on sprout growth, and could justify the hardly perceptible effect of stump height on biomass yield. It implies that stump height represents an important factor for the species’ functional structure and partially confirms our hypothesis 1 that stump height is an important factor for growth and biomass yield in *V. doniana.* The contrasting effect of stump height indicates a modification of resources allocation and net photosynthesis. The fact that basal stem diameter growth rate decreased in the first 45 DAP and then started to increase only at 60 DAP supports that coppicing modifies resources allocation among plant organs. In the first 45 DAP, coppiced trees invested more resources in sprouting and sprouts growth in length than in main stem growth in diameter. Indeed, this should be taken into account in further research aiming at simulating the species’ response to cultivation management, especially to pruning because it modifies architecture, leaf area and consequently photosynthetic processes and crop yield (Marcelis *et al.* 1998, Balandier *et al.* 2000).

In addition, further research should identify the exact distance from which the inhibitory effect of apical buds is suppressed to allow better sprouting and biomass yield. In fact, pruning breaks the inhibitory correlations among tree organs. Removing the top of tree therefore releases the latent buds that develop into new shoots that can be harvested. However, there is a distance over which pruning can suppress this complex inhibitory relationship (Balandier *et al.* 2000). The two stump heights tested in this study suppressed this inhibitory effect and there is need to test low and higher heights.

Further studies are also needed to determine the dynamic of shoot-root ratio in coppiced trees. It is known that coppicing and pruning modify the shoot-root ratio (Balandier *et al.* 2000). The physiological response to coppicing is the modification of tree growth to restore this equilibrium between above and below-ground biomass. This explains our finding that short stumps (higher defoliation) quickly produced higher number of new sprouts to restore this equilibrium. However, the amount of reserve carbohydrates and photosynthetic capacities of subjects determine the speed at which the stumped trees will grow. This differential availability of reserves explains why leafy high stumps grew better and tended to continue producing higher biomass, even with increased pruning frequency. Also, the low suckering capacity observed in coppiced trees indicated that clear-felling as harvesting practice reduces regeneration from rootstock. It implies that this practice which is observed in the natural vegetation (Agossou 2011, N’Danikou *et al.* 2011), will heighten the already existing threat, as fruiting ability is already reduced by leaves harvests.

Findings also indicated that almost all growth parameters increased with initial stem growth. Comparing the effect of initial stem growth as fixed factor and then as random factor, results of the deviance analysis indicated that the effect of IBSD-C cannot be ignored when randomizing the plants in blocks.

Economic studies are also required to evaluate the profitability of this potential plantation crop. As a wild-harvested resource, the majority of farmers will engage in the cultivation of *V. doniana* when there is a clear demonstration of its profitability. In this end, robust environmental economics and crop yield models would be necessary to determine the opportunity cost of *V. doniana* commercial plantation. Furthermore, comparative studies would be desired to determine the profitability of a dual purpose *V. doniana* plantation or agroforestry. In fact, our study revealed a trade-off between leaves and wood production by the coppiced trees. However, a stock of firewood or construction poles can be obtained between two rotation cycles devoted to leaves production.

Finally, biochemical studies are required to compare the nutritional quality of the cultivated versus wild-harvested leaves of *V. doniana.* Apparently the harvested and cooked leaves during the experiment presented similar taste to the wild-harvested, but this should be tested in a formal nutrition protocol.

### 5. Conclusion

The study investigated the effects of coppicing and goat manure application on growth and yield parameters in *V. doniana.* Findings clearly indicated a significant effect of stump height and diameter on production and growth of sprouts, while the effect of goat manure was not prominent. Furthermore, there was a very clear trade-off between production of edible leaves and timber production by the coppiced and periodically pruned trees. Coppicing also reduced formation of root suckers. We concluded that coppicing can be applied to *V. doniana* saplings with 3 cm basal stem diameter or more, and cutting point should not be lower than 40 cm above ground, in order to maximize sprouts growth and biomass yield. The effect of manure on sprouting and biomass yield should be confirmed in further studies. It is also vital to evaluate the effect of treatments on fruiting ability of trees. So far this study is the first of its type in determining the effects of cultivation management on yield of edible biomass, and should serve as background for future investigations towards domestication of *V. doniana* in West Africa.

## Acknowledgements

Authors are grateful to the National Agricultural Research Institute of Benin (INRAB) for providing the land and labour force for the experiment.

## Author’s contributions

SN conceived and designed the study, performed data analysis, wrote and finalized the manuscript. DAT designed and collected data, read and improved the manuscript. COAA and CAH contributed to data gathering, read and improved the manuscript. EGAD gave conceptual advice, read and critically reviewed the manuscript. FAK, RSV and AA gave conceptual advice, read and critically reviewed the manuscript. AA gave final approval of the article. All authors read and approved the final manuscript.

## Funding sources

Funding: This work was supported by New Alliance Trust [Grant # RGNAT04/12].

New Alliance Trust was not involved in study design; in the collection, analysis and interpretation of data; in the writing of the report; and in the decision to submit the article for publication.

## Abbreviations

*CI*: Confidence interval
*DAP*: Days after pruning
*DBH*: Diameter at breast height
*IBSD-C*: Initial basal stem diameter before coppicing
*IBSD-T*: Initial basal stem diameter before transplanting

## References

Achigan-Dako EG, N’Danikou S, Tchokponhoue AD, Assogba-Komlan F, Larwanou L, Vodouhe SR, Ahanchede A. 2014. Sustainable use and conservation of Vitex doniana Sweet: unlocking the propagation ability using stem cuttings. Journal of Agriculture and Environment for International Development, 10843–62. https://dx.doi.org/10.12895/jaeid.20141.195

Agossou OAC. 2011. Pressions anthropiques et stratégies locales de domestication de *Vitex doniana* Sweet dans la commune de Djidja au sud Bénin. Mémoire de Licence. Université d’Abomey-Calav, Faculté des Sciences Agronomiques.

Ahoton LE, Adjakpa JB, Gouda M, Daïnou O, Akpo EL. 2011. Effet des prétraitements de semences du prunier des savanes (vitex doniana sweet) sur la germination et la croissance des plantules. Annales des Sciences Agronomiques, 15.

Akinnifesi FK, Mhango J, Sileshi G, Chilanga T. 2008. Early growth and survival of three miombo woodland indigenous fruit tree species under fertilizer, manure and dry-season irrigation in southern Malawi. Forest ecology and Management, 255546–557. http://dx.doi.org/10.1016/j.foreco.2007.09.025

Atindogbe G, Fonton NH, Fandohan B, Lejeune P. 2012. Caractérisation des plantations privées de teck *(Tectona grandis* Lf) du département de l'Atlantique au Sud-Bénin. Biotechnologie, Agronomie, Société et Environnement, 16441–451.

Balandier P, Lacointe A, Le Roux X, Sinoquet H, Cruiziat P, Le Dizès S. 2000. SIMWAL: a structural-functional model simulating single walnut tree growth in response to climate and pruning. Annals of Forest Science, 57571–585. https://doi.org/10.1093/oxfordjournals.aob.a084308

Berninger F, Nikinmaa E, Sievänen R, Nygren P. 2000. Modelling of reserve carbohydrate dynamics, regrowth and nodulation in a N2-fixing tree managed by periodic prunings. Plant, Cell & Environment, 231025–1040. http://dx.doi.org/10.1046/j.1365-3040.2000.00624.x

Berninger F, Salas E. 2003. Biomass dynamics of *Erythrina lanceolata* as influenced by shoot-pruning intensity in Costa Rica. Agroforestry Systems, 5719–28. http://dx.doi.org/10.1023/A:1022910310082

Davidson R. 1969. Effect of root/leaf temperature differentials on root/shoot ratios in some pasture grasses and clover. Annals of Botany, 33561–569. https://doi.org/10.1093/oxfordjournals.aob.a084308

Dharmakeerthi RS, Senevirathna AMWK, Edirimanne VU, Chandrasiri JAS. 2008. Effect of stock pruning on shoot and root growth of budded polybag plants of Hevea brasiliensis. Natural Rubber Research, 2124–31. http://dx.doi.org/10.1186/2193-1801-1-84

Ferguson JJ. 1994. Growth and yield of bearing and non-bearing citrus trees fertilized with fresh and processed chicken manure. Proceedings of the Florida State Horticultural Society, 10729–32.

Gamal AM, Ragab MA. 2003. Effect of organic manure source and its rate on growth, nutritional status of the trees and productivity of Balady mandarin trees. Assiut Journal of Agricultural Sciences, 34253–264.

Hartmann HT, Kester DE, Davies FT, Geneve RL (Eds.). 2011. Plant propagation: principles and practices, 8th edition. Prentice-Hall, Englewood Cliffs.

Hytönen J. 1994. Effect of cutting season, stump height and harvest damage on coppicing and biomass production of willow and birch. Biomass and Bioenergy, 6349–357. http://dx.doi.org/10.1016/0961-9534(94)E0029-R

Hytönen J, Issakainen J. 2001. Effect of repeated harvesting on biomass production and sprouting of Betula pubescens. Biomass and Bioenergy, 20237–245. http://dx.doi.org/10.1016/S0961-9534(00)00083-0

Kupzow A. 1980. Theoretical basis of the plant domestication. Theoretical and Applied Genetics, 5765–74.

Ky KJM. 2008. *Vitex doniana* Sweet. in Prota 7(1): Timbers/Bois d’oeuvre 1 (Louppe D, Oteng-Amoako AA, Brink M eds.). pp. 578–581. PROTA Foundation, Wageningen, Pays Bas: Backhuys Publishers, Leiden, / CTA, Wageningen:578–581.

Makonda F, Kitojo D, Augustino S, Ruffo C, Ishengoma R, Gillah P, Eriksen S, Msanga H. 2016. *Vitex doniana* Sweet: a potential lesser-known and lesser utilized agroforestry timber species in Kilosa district, Morogoro Tanzania. International Journal of Contemporary Applied Sciences, 3100–113.

Mapongmetsem PM. 2006. Domestication of Vitex madiensis’ in the Adamawa Highlands of Cameroon: Phenology and Propagation. Akdeniz üniversitesi ziraat fakultesi dergisi, 19269–278.

Mapongmetsem PM, Fawa G, Noubissie Tchiagam J-B, Nkongmeneck B-A, Biaou SH, Bellefontaine R. 2016. Vegetative propagation of *Vitex doniana* Sweet from root segments cuttings. Bois et Forêts des Tropiques, 29–37.

Marcelis LFM, Heuvelink E, Goudriaan J. 1998. Modelling biomass production and yield of horticultural crops: a review. Scientia Horticulturae, 7483–111. http://dx.doi.org/10.1016/S0304-4238(98)00083-1

N’Danikou S, Achigan-Dako EG, Tchokponhoue AD, Assogba Komlan F, Vodouhe SR, Ahanchede A. 2015. Improving seedling production for Vitex doniana. Seed Science and Technology, 4310–19. http://doi.org/10.15258/sst.2015.43.1.02

N’Danikou S, Achigan-Dako EG, Tchokponhoué DA, Assogba Komlan F, Gebauer J, Vodouhè RS, Ahanchèdè A. 2014. Enhancing germination and seedling growth in *Vitex doniana* Sweet for horticultural prospects and conservation of genetic resources. FRUITS, 69279–291. http://dx.doi.org/10.1051/fruits/2014017

N’Danikou S, Achigan-Dako EG, Wong JLG. 2011. Eliciting Local Values of Wild Edible Plants in Southern Bénin to Identify Priority Species for Conservation. Economic Botany, 65381–395.

Orozco M, Thienhaus S. 1997. The effect of three rates of chicken manure on the growth and development of cocoa (*Theobroma cacao* L.) trees. Agronomia Mesoamericana, 881–92.

Oumorou M, Sinadouwirou T, Kiki M, Glele Kakaï R, Mensah G, Sinsin B. 2010. Disturbance and population structure of *Vitex doniana* Sweet. in northern Benin, West Africa. International Journal of Biological and Chemical Sciences, 4624–632.

Paukkonen K, Kauppi A. 1998. Effect of coppicing on root system morphology and its significance for subsequent shoot regeneration of birches. Canadian Journal of Forest Research, 281870–1878. 10.1139/cjfr-28-12-1870

Saifuddin M, Hossain ABMS, Osman N, Sattar MA, Moneruzzaman KM, Jahirul MI. 2010. Pruning impacts on shoot-root-growth, biochemical and physiological changes of Bougainvillea glabra. Australian Journal of Crop Science, 4530–537.

Sanoussi A, Ahoton LE, Odjo T. 2012. Propagation of black plum *(Vitex donania* Sweet) using stem and root cuttings in the ecological conditions of South Benin. Tropicultura, 30107–112.

The R Core Team. 2013. R: A language and environment for statistical computing. Vienna: R Foundation for Statistical Computing.

Thomas P, Schiefelbein WJ. 2004. Roles of leaf in regulation of root and shoot growth from single node softwood cuttings of grape (*Vitis vinifera*). Annales of Applied Biology, 14427–37. http://dx.doi.org/10.1111/i.1744-7348.2004.tb00313.x

Tipu SU, Hossain KL, Islam MO, Hossain MA. 2006. Effect of pruning height on shoot biomass yield of Leucaena leucocephala. Asian Journal of Plant Sciences, 51043–1046. https://doi.org/10.3923/ajps.2006.1043.1046

Wilson PJ. 1988. Adventitious rooting in stem cuttings of Eucalyptus grandis Hill ex Maid. University of Natal, Pietermaritzburg,

Wilson PJ. 1994. Contributions of the leaves and axillary shoots to rooting in *Eucalyptus grandis* Hill ex Maid, stem cuttings. Journal of Horticultural Science, 69 999–1007. http://dx.doi.org/10.1080/00221589.1994.11516538

Yasui H, Fujie I. 1971. Studies on the productive structure of Shirakashi (*Cyclobalanopsis myrsinaefolia* Oerst.) coppice-forest managed by selection method. 8. On the growth and biomass of Shirakashi coppice managed by the clear-felling system. Bulletin of the Faculty of Agriculture, Shimane University, 49–55.

Zeder MA. 2015. Core questions in domestication research. Proceedings of the National Academy of Sciences of the United States of America, 1123191–3198. http://dx.doi.org/10.1073/pnas.1501711112

